# Modeling the structure of the frameshift stimulatory pseudoknot in SARS-CoV-2 reveals multiple possible conformers

**DOI:** 10.1101/2020.06.08.141150

**Authors:** Sara Ibrahim Omar, Meng Zhao, Rohith Vedhthaanth Sekar, Sahar Arbabi Moghadam, Jack A. Tuszynski, Michael T. Woodside

## Abstract

The coronavirus causing the COVID-19 pandemic, SARS-CoV-2, uses −1 programmed ribosomal frameshifting (−1 PRF) to control the relative expression of viral proteins. As modulating −1 PRF can inhibit viral replication, the RNA pseudoknot stimulating −1 PRF may be a fruitful target for therapeutics treating COVID-19. We modeled the unusual 3-stem structure of the stimulatory pseudoknot of SARS-CoV-2 computationally, using multiple blind structural prediction tools followed by μs-long molecular dynamics simulations. The results were compared for consistency with nuclease-protection assays and single-molecule force spectroscopy measurements of the SARS-CoV-1 pseudoknot, to determine the most likely conformations. We found several possible conformations for the SARS-CoV-2 pseudoknot, all having an extended stem 3 but with different packing of stems 1 and 2. Several conformations featured rarely-seen threading of a single strand through the junction formed between two helices. These structural models may help interpret future experiments and support efforts to discover ligands inhibiting −1 PRF in SARS-CoV-2.

## INTRODUCTION

The COVID-19 pandemic caused by the novel Severe Acute Respiratory Syndrome coronavirus 2 (SARS-CoV-2) has spread across the globe since the virus emerged in late 2019 (1). Given the high infectivity of SARS-CoV-2 and the novel immunological challenge it poses to human hosts, epidemiological modeling suggests that recurring outbreaks with elevated mortality can be expected even despite successful public-health responses, until vaccines or preventive drugs can be found to inhibit transmission (2). The discovery of effective treatment therapeutics is thus one of the central goals of research into COVID-19 (3).

One potential target for treatment is the frameshift-stimulatory pseudoknot found between the overlapping ORF1a and ORF1b in the SARS-CoV-2 genome (4). Like other human coronaviruses (5), SARS-CoV-2 depends on −1 programmed ribosomal frameshifting (−1 PRF) to produce essential proteins at regulated levels (6). In −1 PRF, a shift in reading frame is stimulated at a specific location in the RNA message by a structure in the mRNA—typically a pseudoknot—located 5–7 nt downstream of a ‘slippery’ sequence, thereby generating more than 1 protein from the same message (7, 8). The ratio of frameshifted gene products must often be held within a tight range for optimal propagation of the virus, hence disrupting −1 PRF by modulating the efficiency of frameshifting can attenuate the virus. Indeed, inhibiting −1 PRF was found to suppress replication of the close relative SARS-CoV-1 by orders of magnitude (9, 10), suggesting that the same strategy may be effective against SARS-CoV-2.

The structure of the frameshift-stimulatory pseudoknot has not yet been solved for either SARS-CoV-1 or SARS-CoV-2, however, hindering structure-based drug-discovery efforts. The primary sequences of these two pseudoknots are almost identical, differing by a single nucleotide, hence their secondary structure is expected to be the same. Evidence from computational methods, nuclease protection assays, and 2D NMR spectroscopy applied to the SARS-CoV-1 pseudoknot (11, 12) indicates it has an unusual 3-stem architecture: whereas pseudoknots typically consist of 2 interleaved stems and loops, here the second loop is greatly extended and a third stem-loop combination forms within it (Fig. 1). Bulged adenine residues in stem 2 (S2) and stem 3 (S3) seem to play important functional roles, as mutating them to cytosine abolished or reduced −1 PRF (respectively for the bulges in S2 and S3) (12). However, the full 3D structure has never been solved for any 3-stem pseudoknot.

**Figure 1:**
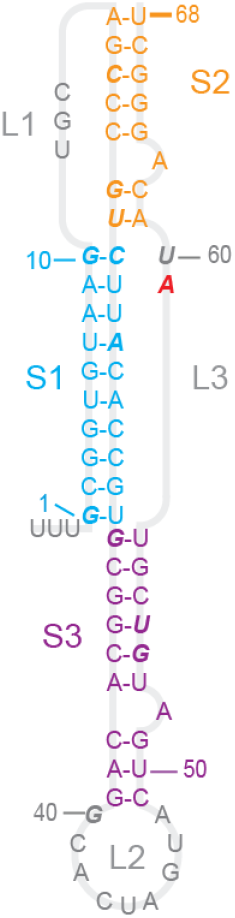
SARS-CoV-2 pseudoknot primary and secondary structure. The sequence is color-coded by secondary structure (S1: cyan, S2: orange, S3: purple, loops: grey). The only difference from SARS-CoV-1 is that A59 (red) is changed to C59 in the latter. Bases shown in italic are protected against nuclease digestion in SARS-CoV-1.

Computational modeling provides an alternative approach to characterizing the structure of these pseudoknots, but such modeling has been limited to date. One study assembled a 3D structure of the SARS-CoV-1 pseudoknot by hand with the Sybyl chemical modeling package before equilibrating briefly with 1 ns of molecular dynamics (MD) simulation (13), and another study used the Rosetta FARFAR2 platform (14, 15) to make a ‘blind’ prediction of the structure of the SARS-CoV-2 pseudoknot (16). Here, we have modeled the structure of the SARS-CoV-2 pseudoknot more extensively, using blind predictions from multiple platforms as inputs for μs-long MD simulations to examine the stability of the structures. We also assessed the ensemble of structures observed in the simulations for their consistency with previous work on the biochemical and biophysical properties of the SARS-CoV-1 pseudoknot to identify the most likely structural models. We found several possibilities, all sharing an extended S3 helix but differing in the S1/S2 packing and junction with S3.

## METHODS

### Blind structure prediction

Initial structures for input into MD simulations were obtained using multiple platforms for blind RNA structure prediction: SimRNA (17), Rosetta FARFAR2 (14, 15), RNAComposer (18), RNAvista (19), RNA2D3D (20), and Vfold (21). For the blind predictions, we assumed the secondary structure shown in Fig. 1, based on previous characterization of the secondary structure of the SARS-CoV-1 pseudoknot (12).

### MD simulations

Models from blind structure predictions were used as starting structures for all-atom MD simulations in explicit solvent. The models were protonated at pH 7 using Molecular Operating Environment software. The pseudoknots were parameterized using the f99bsc0_chiOL3 force-field and were solvated in optimal point charge water boxes of 12 Å using the tleap module of Amber18 (22). The solvated systems were first neutralized using sodium ions, then their salinities were adjusted to 0.15 M NaCl. Each model was simulated under two conditions: without Mg^2+^ ions, or with six Mg^2+^ ions placed initially at the junction between S1 and S3 as well as along the backbone of S2. The solvated systems were energy-minimized then heated to 310K with heavy restraints of 10 kcal/mol/A^2^ on the backbone phosphate atoms. These restraints were gradually removed and the unrestrained systems were then simulated for 1 μs at constant pressure.

### Analysis of simulated models

Analysis was performed using CPPTRAJ of AmberTools. Different conformations of the pseudoknot within each simulation were clustered based on the root-mean-squared deviation (RMSD) of residues G1–G40 and C49–G66 (omitting the residues in L2 and at the 3′ terminus, which tended to have large fluctuations), using the hierarchical agglomerative approach. The representative structures of the three most populated clusters of each model were visually assessed for helical distortions in S1, S2 and S3. The hydrogen bonding of the residues identified as protected in SARS-CoV-1 by nuclease-protection assays (12) as well as of bulged adenines and residues in L1 were also calculated, reporting the interactions formed by hydrogen bond donors and acceptors between two bases or between a base and the backbone atoms of other residues that were present for at least the last 100 ns of the trajectory. The root-mean-squared fluctuation (RMSF) of each residue was also calculated.

## RESULTS

We made initial estimates of the pseudoknot structure using a variety of tools that have been developed for blind prediction of RNA structure (14, 15, 17–21), where the only input was the sequence and the expected secondary structure, as shown in Fig. 1. Some of these predictions were rejected as being implausible (*e.g*. containing topological knots, which cannot occur) or being obviously inconsistent with the nuclease protection data. The remainder are illustrated in Fig. 2; note that those in Fig. 2F–H were also reported previously in a separate study (16). These predictions were quite varied. They all showed S3 as an extended helix lacking obvious contacts with the rest of pseudoknot except right near the junction with S1. However, the arrangement of S1 and S2 and L1 and L3 differed significantly between the predictions. In several models (Fig. 2A–D), S1 and S2 were packed very tightly, leading to distorted or broken base-pairs in order to accommodate the packing. A couple of models (Fig. 2E, G) showed an unusual quasi-knotted structure where the 5′ end was threaded through the junction between S1 and S3, a fold topology that has only been seen previously in exoribonuclease-resistant RNAs (23). Another model (Fig. 2H) showed a similar fold topology, but this time with L3 (near the 3′ end) threaded through the junction between S1 and S2. Note that in several cases, the first three nucleotides upstream of the 5′ end of the pseudoknot (UUU) were included in the modeling, in order to distinguish the possibilities for 5′-end threading.

**Figure 2:**
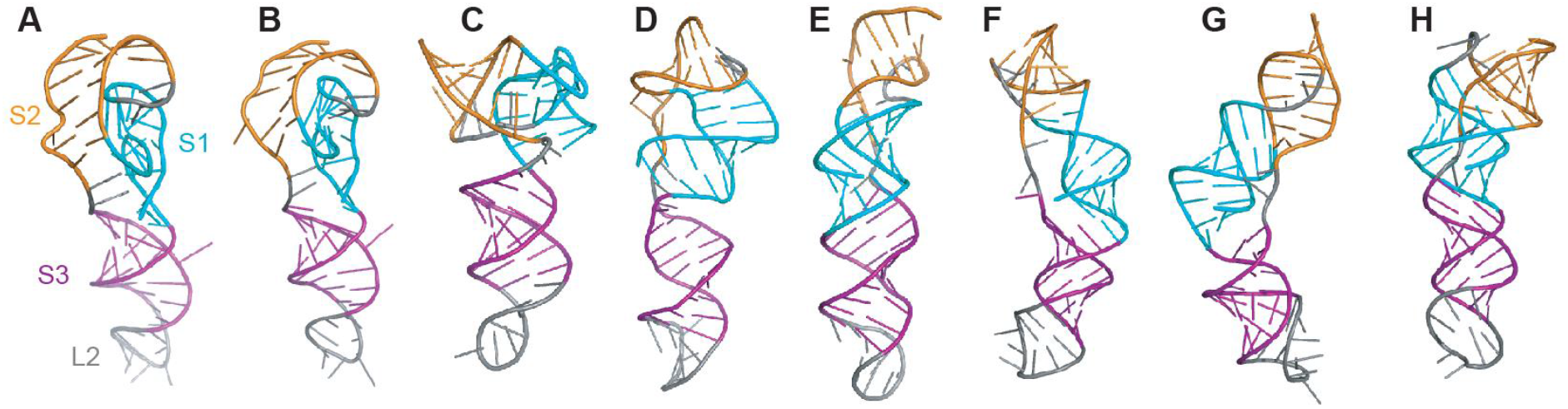
Blind predictions of pseudoknot structure. In each case, secondary structure is color-coded (S1: cyan, S2: orange, S3: purple, loops: grey).

To examine if the blind predictions were dynamically stable, we used them to initiate extended all-atom molecular dynamics simulations. Each structure in Fig. 2 was simulated for at least 1 μs in explicit solvent under two conditions: with NaCl only, or with both NaCl and Mg^2+^ ions. Both conditions were used because not all pseudoknots require Mg^2+^ ions to fold (24, 25), and it is unclear if Mg^2+^ ions are essential for the SARS-CoV-2 pseudoknot. The first part of the simulation was treated as an equilibration phase and only the last 500 ns of the simulation was examined in each case. Because the simulations were dynamic, we clustered the structures occupied in the simulations by RMSD and examined the centroid (representative) structures of the three most occupied clusters.

The initial models that involved the tightest packing of S1 and S2 (Fig. 2A–D) led, after equilibration in MD simulations, to structures that featured various combinations of significant defects in the expected base-pairing for S1, defects in S2, and/or a lack of tertiary contacts (Fig. S1). In some cases, these structures were sufficiently unstable that they unfolded substantially. This set of models was therefore rejected as very unlikely to be correct. The other initial models yielded structures under at least one of the MD simulation conditions that were more plausible, and they were thus analyzed in more detail. The results could be arranged into three groups: structures without any threading at either end (Fig. 3), structures with the 5′ end threaded through the S1/S3 junction (Fig. 4), and a structure with L3 threaded through the S1/S2 junction (Fig. 5).

**Figure 3:**
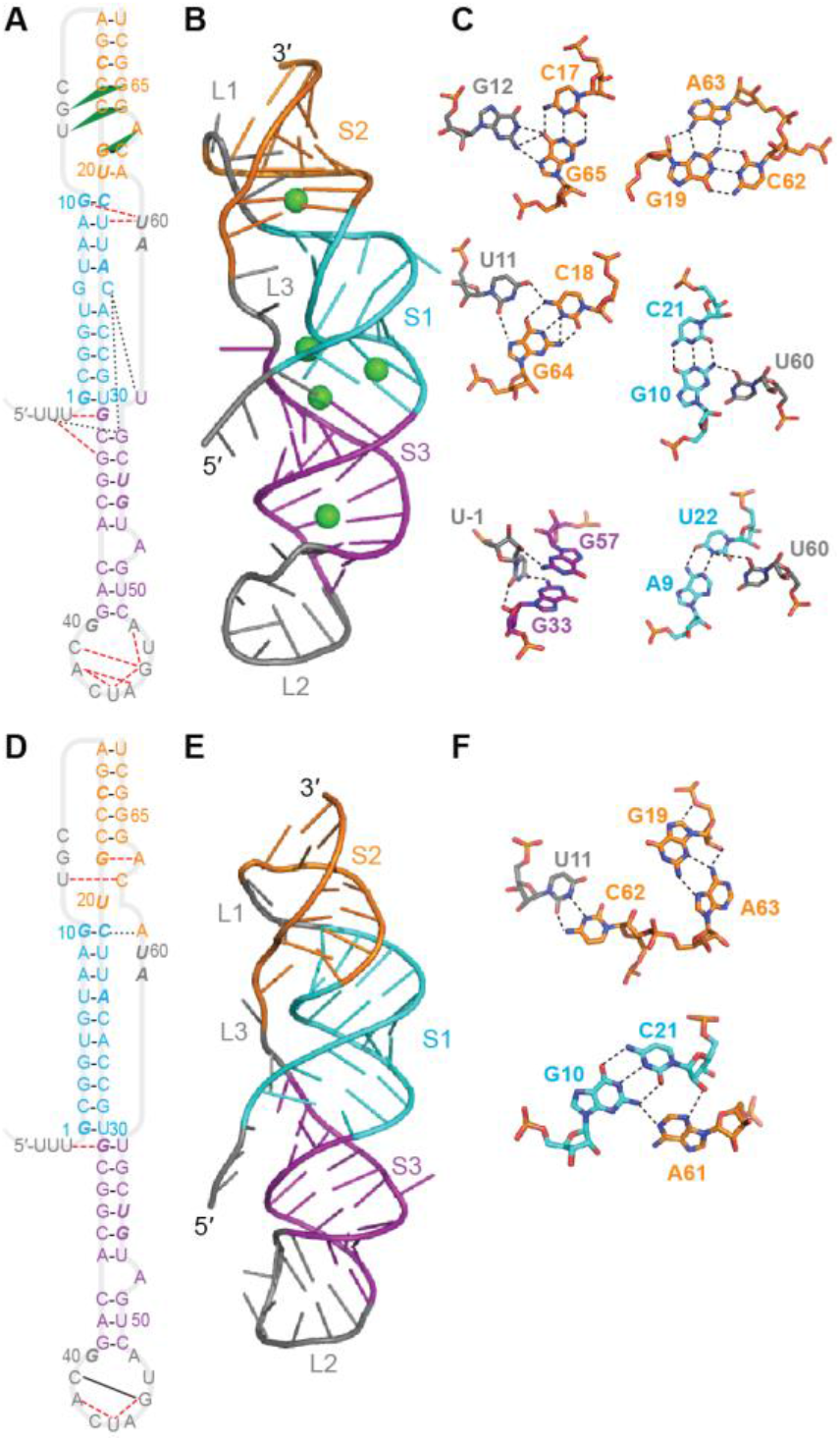
Representative structures from MD simulations of unthreaded model. (A–C) Structure from simulation of Fig. 2F with Mg^2+^. (A) Secondary structure with tertiary contacts. Solid lines: canonical base-pairs; dashed lines: non-canonical base-base hydrogen bonds; dotted lines: base-backbone hydrogen bonds; green triangles: triples or triple-like interactions. Bases shown in italic are protected against nuclease digestion in SARS-CoV-1. (B) Representative 3D structure of most populated cluster. Green spheres: Mg^2+^ ions. (C) Close-up view of key tertiary contacts. (D–F) Structure from simulation of Fig. 2F without Mg^2+^. (D) There are fewer tertiary contacts than with Mg^2+^. (E) Representative 3D structure of most populated cluster. (F) Close-up view of key tertiary contacts.

**Figure 4:**
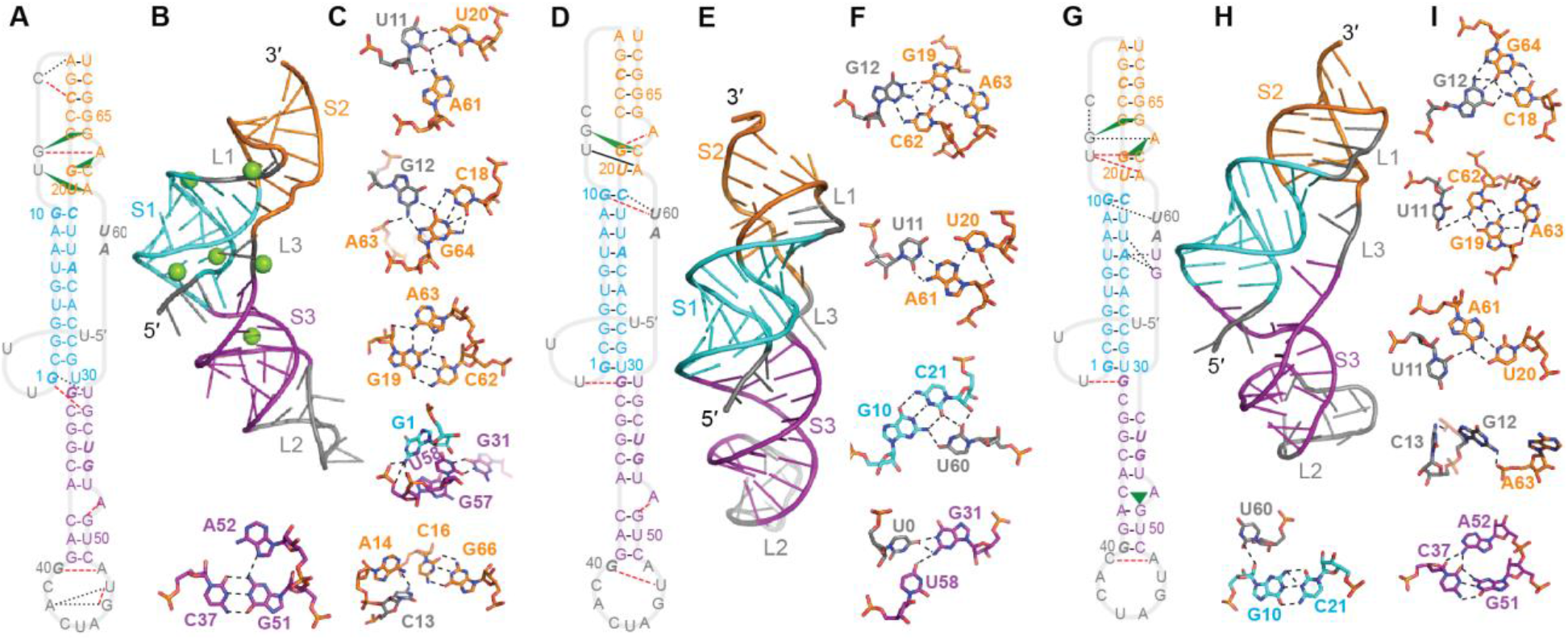
Representative structures from MD simulations of models with 5′-end threading. (A–C) Structure from simulation of Fig. 2G with Mg^2+^. (A) Secondary structure with tertiary contacts. Solid lines: canonical base-pairs; dashed lines: non-canonical base-base hydrogen bonds; dotted lines: base-backbone hydrogen bonds; green triangles: triples or triple-like interactions. Bases shown in italic are protected against nuclease digestion in SARS-CoV-1. (B) Representative 3D structure of most populated cluster. Mg^2+^ ions shown in green. (C) Close-up view of key tertiary contacts. (D–F) Structure from simulation of Fig. 2E without Mg^2+^. (G–I) Structure from simulation of Fig. 2G without Mg^2+^ showing opening of the top of S3.

**Figure 5:**
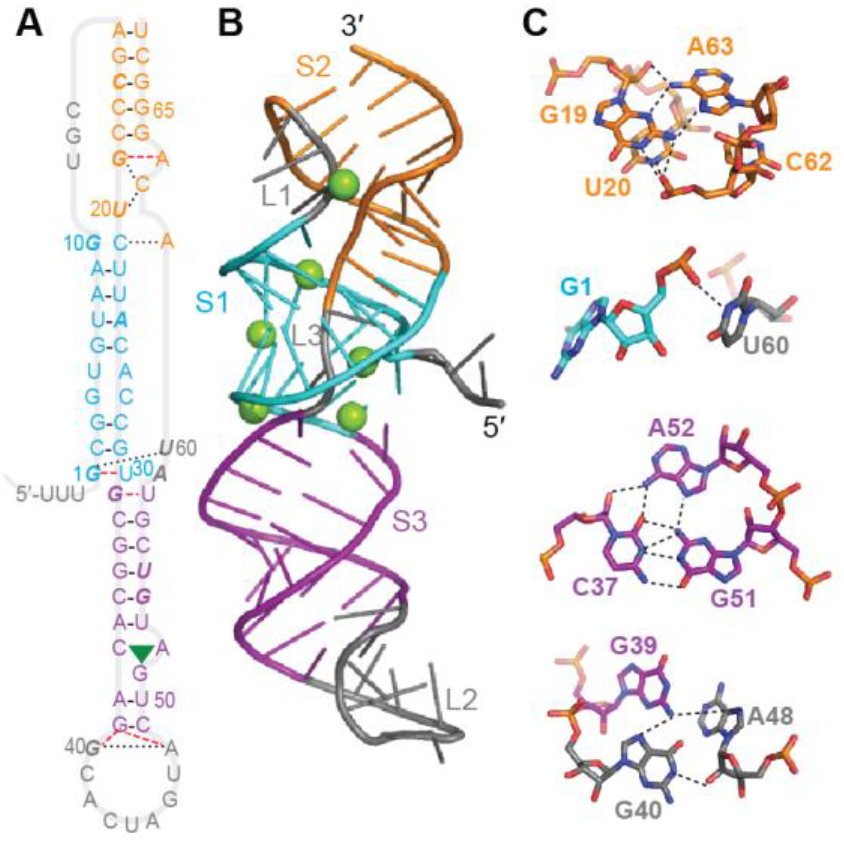
Representative structures from MD simulations of models with L3 threading. (A–C) Structure from simulation of Fig. 2H with Mg^2+^. (A) Secondary structure with tertiary contacts. Solid lines: canonical base-pairs; dashed lines: non-canonical base-base hydrogen bonds; dotted lines: base-backbone hydrogen bonds; green triangles: triples or triple-like interactions. Bases shown in italic are protected against nuclease digestion in SARS-CoV-1. (B) Representative 3D structure of most populated cluster. Mg^2+^ ions shown in green. (C) Close-up view of key tertiary contacts.

Considering first the structures that were more similar to standard H-type pseudoknots, without any threading of either end through stem junctions, the representative structures of the most populated clusters with and without Mg^2+^ are shown in Fig. 3. With Mg^2+^ (Fig. 3A–C), several triples and triple-like interactions were identified in S2/L1 (Fig. 3A, triangles), and U60 in L3 was seen to bond with both G10 and U22 in S1. Base-pairs in S1 (G6:C25) and S3 (G31:U58) were disrupted, in favor of base-backbone bonding (Fig. 3A, dotted lines) between C25 and both G57 and U58, and a wobble pair with the end of the linker (G31:U0). A network of base-base hydrogen-bond interactions (Fig. 3A, dashed lines) was also seen in L2. The stabilization from the hydrogen bond networks led to low fluctuations, even in L2, and the RNA spent almost of its time in the top 3 clusters (Fig S2A). For the simulations without Mg^2+^ (Fig. 3D–F), the disrupted base-pairs in S1 and S3 were restored, but the triples in S2/L1 were absent and the pairing in the lower part of S2 was disrupted in favor of non-canonical interactions between U11:C62 and G19:A63. However, the sparser hydrogen bond network led to higher fluctuations and a greater diversity of clusters than with Mg^2+^ present (Fig. S2B).

Turning next to the representative structures of the most populated clusters with the 5′ end threaded through the S1/S3 junction, we found several possibilities. With Mg^2+^ present (Fig. 4A–C), again several triples/triple-like interactions were identified in S2/L1, two of them the same as in Fig. 3A–C. The opening G:U pair in S1 was disrupted in favor of interactions with G57 and U58, to accommodate the threading of the 5′ end through the S1/S3 junction, and S3 was extended by a non-canonical pair between G40 and A48. L2 was again structured by a network of hydrogen bonds, but L3 did not interact with any other part of the structure other than a coordinated Mg^2+^ ion. Without Mg^2+^, two different structures were seen. In the first (Fig. 4D–F), the 5′ end was threaded through the S1/S3 junction without disrupting the base-pairing near the junction, stabilized by bonding between G31 and U0. Only one triple was seen in S2/L1, but it was also bonded with the bulged A63 in S2. The U20:A61 pair at one end of S2 was distorted to include interactions between U11 and A61, and U60 in L3 was bonded to both G10 and C21, analogous to the situation in Fig. 3A–C. L2 was partially structured by interactions between G40 and U47. In the second structure without Mg^2+^ (Fig. 4G–I), the first two base-pairs in S3 next to the S1/S3 junction were opened to facilitate the 5′-end threading, stabilized by bonding between U0 and G31, with the unpaired G57 and U58 forming H-bonds with the backbone at A24 and U23. Two of the same triples/triple-like interactions were seen as in Figs. 3A–C and 4A–C, and once again U60 in L3 interacted with G10 in S1 (although this time via a base-backbone bond). The bottom of S3 was also re-arranged, forming a triple with the bulged A52 and extending the stem with the pair G40:C49 and non-canonical bonding between C41 and A48. As above, the simulation with Mg^2+^ showed smaller fluctuations than those without Mg^2+^; the fraction of the trajectory spent in the top 3 clusters was very high (over 90%) for the simulation with Mg^2+^ and the first model without Mg^2+^, but lower (66%) for the second model without Mg^2+^ (Fig S3).

Lastly, we considered the representative structure of the most populated cluster with L3 threaded through the S1/S2 junction. In Fig. 5A–C, only the result with Mg^2+^ is shown, as the structure was unstable without Mg^2+^. Here, the threading of L3 disrupted base-pairing in the bottom of S2 and the middle of S1, although the two strands of S2 continued to interact via base-backbone hydrogen bonds and S1 still retained a fairly regular helical structure; U60 bonded with the backbone of G1 to help stabilize the threading of L3. In S3, the same triple formed as in Fig. 4G–I, but without the reconfiguration and extension of the stem. L2 was partially structured by base-backbone bonds between G40, A48, and the closing base-pair of S3. The top three clusters comprised 78% of the MD trajectory, with relatively low fluctuations (Fig. S4).

## DISCUSSION

Perhaps the most surprising aspect of this work is that we find three very different fold topologies that are stable and persistent in long MD simulations. Such a result contrasts with previous work simulating the pseudoknot from SARS-CoV-1, which reported only a single structure that was somewhat similar to the models with 5′-end threading, although the additional nucleotides completing the threading were not included in the model (13). This difference can be explained by the fact that only a single initial structure was explored in that work: the three fold topologies we observe are sufficiently different that they cannot interconvert without substantial unfolding of the S1/S2 region, and they are each sufficiently stable that such unfolding is very unlikely (as seen in our simulations). Furthermore, the existence of multiple structures has been seen previously in various frameshift signals (26–28). Indeed, it is entirely consistent with the hypothesis (29–31) that high-efficiency stimulatory structures such as that from SARS-CoV-2—which induces −1 PRF at a rate of ~20–30% (6)—have high conformational heterogeneity and hence forms more than one structure.

Each of the models presented above is generally consistent with previous experimental characterizations of the SARS-CoV-1 pseudoknot. Single-molecule force spectroscopy of pseudoknot unfolding for SARS-CoV-1 found a broad unfolding force distribution with high forces, indicating the presence of significant tertiary contact formation (29). Consistent with this observation, all of the models feature tertiary contacts and hydrogen-bond networks stabilizing the 3D structure, although the unthreaded model without Mg^2+^ lacked the triples found in the other models. Turning to the results from nuclease-protection and mutation experiments (12), each model contained features that could be matched to protected residues, such as triples, hydrogen-bond networks, or steric protection; the participation of A63 in S2 in triples was also consistent with work showing that mutating A63 can dramatically lower the −1 PRF efficiency. Again, the unthreaded model without Mg^2+^ was most lacking in these features, suggesting that it is the model that is least consistent with the experimental data. We note, however, that none of the structural models provided an obvious explanation for a few of the protected nucleotides, such as G54 and U55.

It is still unknown if Mg^2+^ is essential for the folding of this pseudoknot. Our modelling, however, suggests that Mg^2+^ helps to stabilize the structures. In every case, the fluctuations were reduced with Mg^2+^ present, and in most cases the presence of Mg^2+^ stimulated a denser network of hydrogen bonds. The role of Mg^2+^ was particularly important for threading L3 through the S1/S2 junction (Fig 5): Mg^2+^ was found to be essential for maintaining the integrity of S1, as the tight packing of the backbone that was needed could not be accommodated without the countervailing charge from the ions.

The threaded fold topologies are particularly interesting: although they have been observed in exoribonuclease-resistant RNAs, where the 5′ end is threaded through a ring closed by a pseudoknot (23), no such fold has been seen in a frameshift-stimulatory structure. We note that threading of either the 5′ end (as in Fig. 4) or L3 (as in Fig. 5) requires that the different parts of the structure fold in a specific order. For example, S2 would likely need to form last for 5′-end threading, else the RNA upstream of the pseudoknot would be unable to thread through the S1/S3 junction, whereas S1 would likely need to form last for L3 threading, else the downstream RNA would be unable to thread through the S1/S2 junction. As a result, the pseudoknot would be expected to populate multiple conformers, dictated by the kinetic partitioning between the different possible pathways (32).

The structural models described above will be helpful for future experimental analyses of the SARS-CoV-2 pseudoknot. X-ray scattering profiles can be predicted from these models and used to analyze small- and wide-angle x-ray scattering measurements, to confirm which (if any) of these conformations are populated and in what kind of mixture (33, 34). The models could also be compared to single-molecule measurements of pseudoknot folding, which could detect heterogeneous populations of different conformers and characterize the sequence of intermediate states formed during the folding of each one (28, 35, 36). These models should also prove useful for drug discovery efforts, facilitating structure-based searches for compounds that attenuate the virus by altering −1 PRF.

## Acknowledgements

We thank Compute Canada for providing exceptional access to computational resources for this project. This work was supported by the Canadian Institutes for Health Research and Alberta Innovates.

