## Supporting Figures 1-4 for "Modeling the structure of the frameshift stimulatory pseudoknot in SARS-CoV-2 reveals multiple possible conformers"

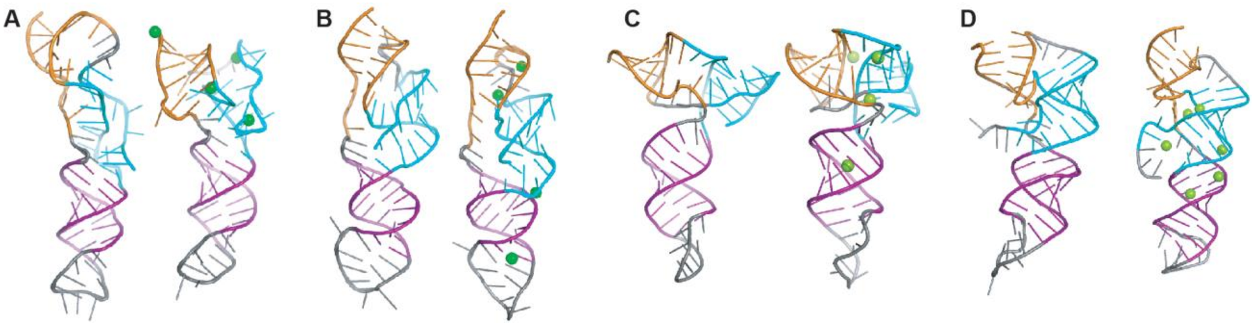


**Figure S1: Selected structures from simulations with significant secondary structure disruption**. Representative structures of the most populated cluster from simulations of (A) Fig. 2A, (B) Fig 2B, (C) Fig. 2C, and (D) Fig. 2D show significant disruption of the secondary structure. In each panel, the figure on the right is from simulations with $Mg^{2+}$, that on the left is from simulations without $Mg^{2+}$.

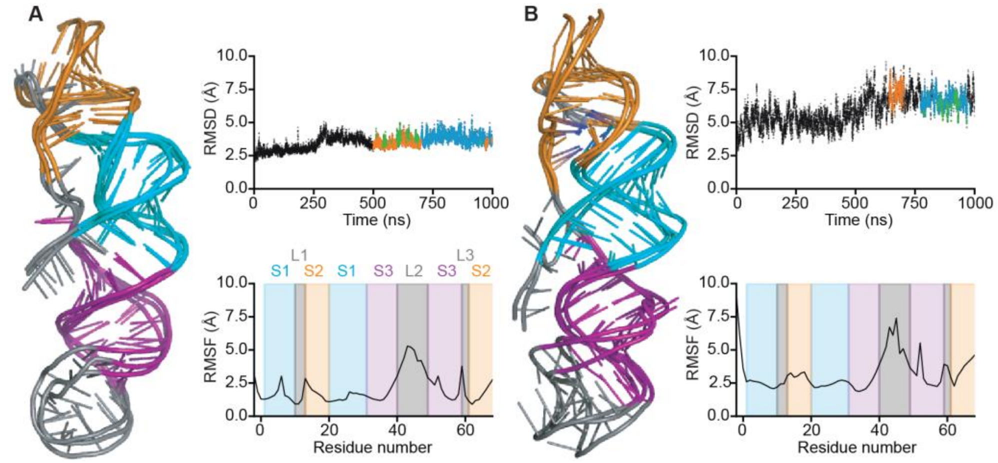


**Figure S2: MD simulations of unthreaded model**. (A) Overlay of the 3D structure of the 3 most populated clusters from simulations of Fig. 2F with $Mg^{2+}$ (ions not shown for clarity). Top inset: RMSD vs time, showing when each of the 3 most populated clusters was populated during the last 500 ns of the simulation (blue: top cluster, orange: second cluster, green: third cluster). Bottom inset: RMSF for each residue. (B) The same for simulations without $Mg^{2+}$.

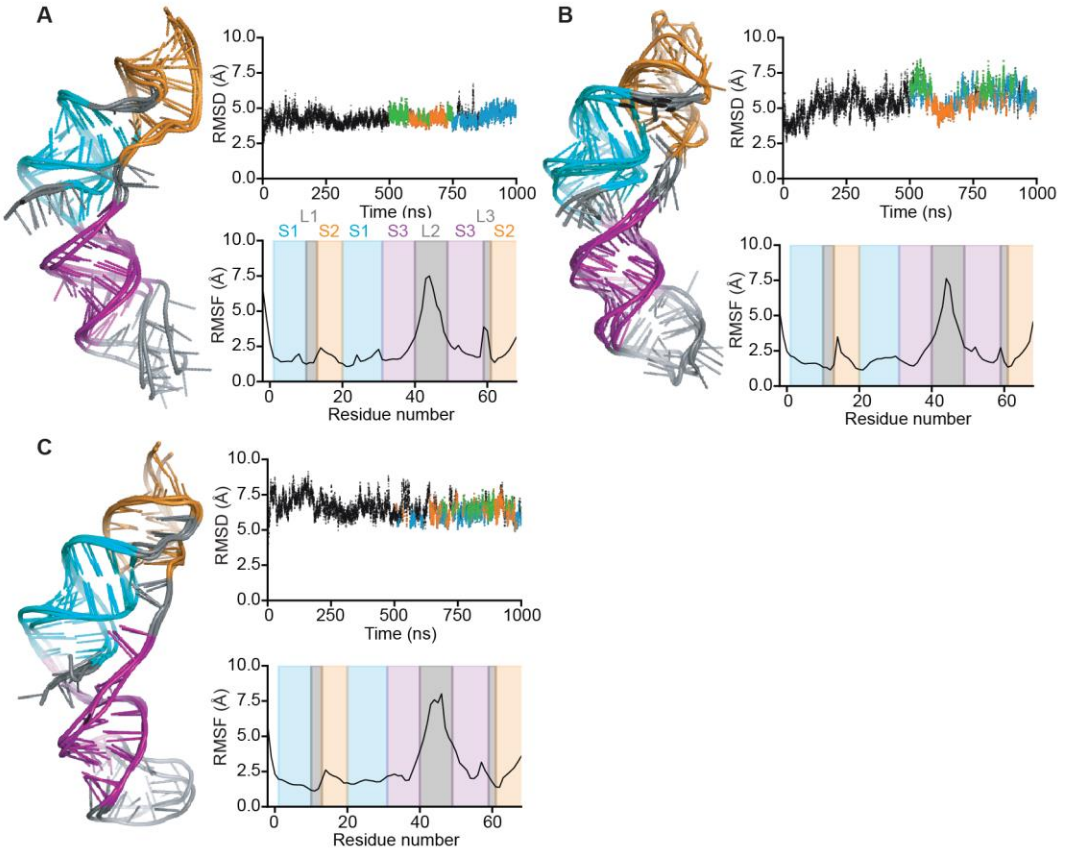


**Figure S3: MD simulations of models with 5′-end threading**. (A) Overlay of the 3D structure of the 3 most populated clusters from simulations of Fig. 2F with $Mg^{2+}$ (ions not shown for clarity). Top inset: RMSD vs time, showing when each of the 3 most populated clusters was populated during the last 500 ns of the simulation (blue: top cluster, orange: second cluster, green: third cluster). Bottom inset: RMSF for each residue. (B) The same for simulations of Fig. 2E without $Mg^{2+}$. (C) The same for simulations of Fig. 2G without $Mg^{2+}$.

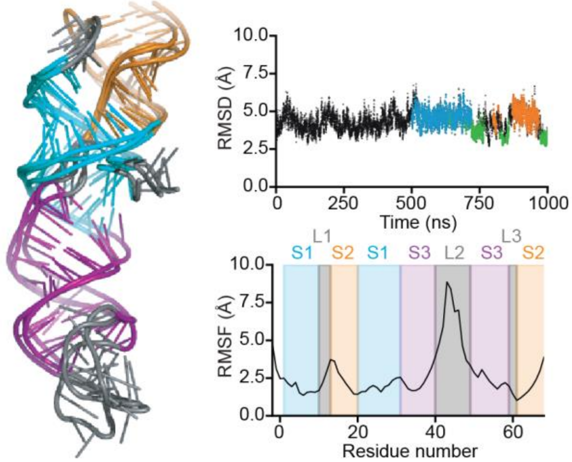


**Figure S4: MD simulations of model with L3 threading**. (A) Overlay of the 3D structure of the 3 most populated clusters from simulations of Fig. 2H with $Mg^{2+}$ (ions not shown for clarity). Top inset: RMSD vs time, showing when each of the 3 most populated clusters was populated during the last 500 ns of the simulation (blue: top cluster, orange: second cluster, green: third cluster). Bottom inset: RMSF for each residue.
